# Sepsis-induced brain state instability

**DOI:** 10.1101/2022.03.09.482831

**Authors:** Annu Kala, Susan Leemburg, Karel Jezek

## Abstract

Sepsis-associated brain dysfunction (SABD) is a frequent severe complication of sepsis and the systemic inflammatory response syndrome. It is associated with high mortality and a majority of survivors suffer long-term neurological consequences. Here, we aimed at better understanding the effects of SABD on oscillatory brain states in an acute rat model of sepsis induced by high dose LPS (10 mg/kg). To focus on intrinsically generated brain state dynamics, we used a urethane model that spares oscillatory activity in REM- and NREM-like sleep states. Soon after the intraperitoneal LPS injection we observed a robust instability of both oscillatory states resulting in tripling amount of state transitions that lasted for several hours, although the overall time spent in either state did not change. Analysis of power spectra showed opposing shifts in low frequency oscillations (1-9 Hz) that resulted in increased similarity between both states in 2-D state space. The described spectral characteristics of sepsis-induced EEG state instability might point to a mechanism underlying severe sleep fragmentation as described both in sepsis patients and in SABD animal models.

## 1. Introduction

Sepsis, a state of dysregulated host response against an infection, affects around 30 million people each year and has 33% mortality (Annane and Sharshar, 2015; Rhee et al., 2017). The brain is among the first impacted organ systems: up to 70% of sepsis patients suffer sepsis-associated brain dysfunction (SABD) early during the disease (Czempik et al., 2020). Out of those who survive, 20 - 25% experience post-sepsis syndrome, which, besides other organ systems, often affects the brain as well (Annane and Sharshar, 2015; Chavan et al., 2012), with a wide spectrum of signs like sleep disturbances, concentration impairment, cognitive deterioration or psychiatric conditions.

Dysregulated sleep in its various forms is one of the dominant neurological signs in intensive care unit patients including the acute stage of sepsis (Gofton and Young, 2012; Richards and Bairnsfather, 1988). Sleep in septic patients is often markedly discontinuous and interrupted by frequent awakening (Boyko et al., 2017; Weinhouse and Schwab, 2006). Even in cases where the cumulative sleep time might not be reduced, the duration of continuous sleep periods is severely shortened, impairing sleep quality and the physiological functions of the sleep related machinery (Dzierzewski et al., 2020; Opp, 2005; Tononi and Cirelli, 2014; Zamore and Veasey, 2022).

This sleep fragmentation has been described in humans as well as in animal models of generalized inflammation or sepsis (Baracchi et al., 2011; Weinhouse and Schwab, 2006). Fragmented sleep is considered to be an important factor in the development of other neurological signs in both acute systemic inflammation and in chronic post-sepsis syndrome (Song et al., 2021). While in the acute phase it promotes further patient’s deterioration across multiple systems among others by preventing efficient tissue restoration or by deepening the immune dysfunction, in the long term, as part of post-sepsis syndrome sleep fragmentation may be linked to cognitive functions, mental stability and fatigue (Song et al., 2021; Sonneville et al., 2013; Spira et al., 2014)

Besides disruptions in sleep/wake rhythmicity, sepsis patients suffer from a variety of pathological brain activity patterns, including electrographic seizures and periodic discharges, as well as other changes in spectral and oscillatory activity. The rate of mortality and severe disability were found to be directly proportional to the occurrence of electrographic seizures and periodic discharges, stressing the importance of early monitoring of brain activity (Oddo et al., 2009). Additionally, increased slow wave activity and decreased alpha power in the EEG were associated with mortality in the ICU and predicted the occurrence of delirium in septic shock patients (Azabou et al., 2015; Hosokawa et al., 2014; Kinoshita et al., 2021).

These findings were partly confirmed in animal SABD models. In a rat caecal-ligation-and-puncture model, sepsis led to acute sleep fragmentation and suppression of REM sleep, as well as to an increase in the overall amount of dark phase NREM sleep (Baracchi et al., 2011). Interestingly, slow wave activity was reduced during these periods of increased NREM sleep. A different study in anaesthetized rats using LPS-induced sepsis found reduced power in the 8 – 13 Hz range, with no effects on delta power (Semmler et al., 2008). Under non-septic conditions, immunological challenges such as low doses of LPS modulate sleep as well (Opp, 2005). Low doses of LPS caused NREM fragmentation in rats and increased NREM slow wave activity while suppressing REM sleep (Lancel et al., 1995). Some of the observed effects of LPS on EEG spectra were similar to those of sleep deprivation, which might signify a common mechanism of EEG regulation in both conditions. However, NREM fragmentation was specific to LPS administration, and typically does not occur during recovery after sleep deprivation (Borbély et al., 1984; Trachsel et al., 1986). Similar effects of mild inflammation caused by LPS or Lipid A on NREM and REM sleep have been observed in rabbits (Krueger et al., 1986). Other studies in rats confirmed that low doses of LPS cause NREM discontinuity, but showed reduced NREM slow wave activity instead (Kapás et al., 1998)

Apart from affecting overall sleep architecture, LPS has region-specific effects on brain oscillatory activity: (Mamad et al., 2018) observed an acute slowing of hippocampal theta, with a concurrent increase in hippocampal delta frequency and power in awake rats. By contrast, prefrontal cortex showed no change in delta frequency, but exhibited reductions in theta frequencies and power (Mamad et al., 2018). The varying consequences of LPS on delta power in NREM could therefore be a result of species- or dose-specific effects, as well as time of injection and recorded brain region.

Electrophysiological changes of brain activity in sleep thus could serve as a potential early biomarker of sepsis and its related outcomes. Here, we aim to understand the dynamics of REM-like and NREM-like states and their transitions during severe acute systemic inflammation caused by a high dose of LPS (10 mg/kg). We used urethane anaesthesia as a model which mimics the unconscious state of sleep not interrupted by awakening, and produces sleep-like REM and NREM brain activity patterns (Clement et al., 2008). Local field potentials from the CA3 region of the hippocampus were recorded in saline-injected and LPS-injected rats. LPS injection resulted in profound shortening of REM and NREM states. Additionally, spectral similarity between REM and NREM-like states increased, particularly in lower frequencies (1-9 Hz). The observed spectral changes leading to higher brain states similarity might be an important contributing factor to their instability and thus to sepsis-related sleep disturbances and associated deficits in cognitive functions.

## 2. Materials and Methods

### 2.1. Animals

Twelve adult Long-Evans rats weighing 400-500 grams were used. All protocols were approved by the Ethical Committee of the Ministry of Education, Youth and Sports of the Czech Republic (approval no. MSMT-12084/2019) according to the Guide for the Care and Use of Laboratory Animals (Protection of Animals from Cruelty, Law Act No. 246/92, Czech Republic). The animals were housed individually in transparent polycarbonate cages and were kept in a 12:12 light/dark cycle with food and water *ad libitum*. Experiments were carried out during the light phase.

### 2.2. Experimental Setup

To investigate the effects of acute systemic inflammation, two groups of rats were used (LPS and CTRL, N = 6 each, Fig. 1A). One hour after induction of urethane anesthesia, rats in both groups were injected with sterile saline (2ml/kg, i.p.). Then, baseline hippocampal activity was recorded for three hours. After this, rats in the LPS group were injected intraperitoneally with 10 mg/kg lipopolysaccharide and were recorded for another three hours. Rats in the CTRL group received a second saline injection instead. After the second recording period, blood serum samples and brains were collected for IL-1β quantification and histological verification of electrode placement.

**Fig. 1.**
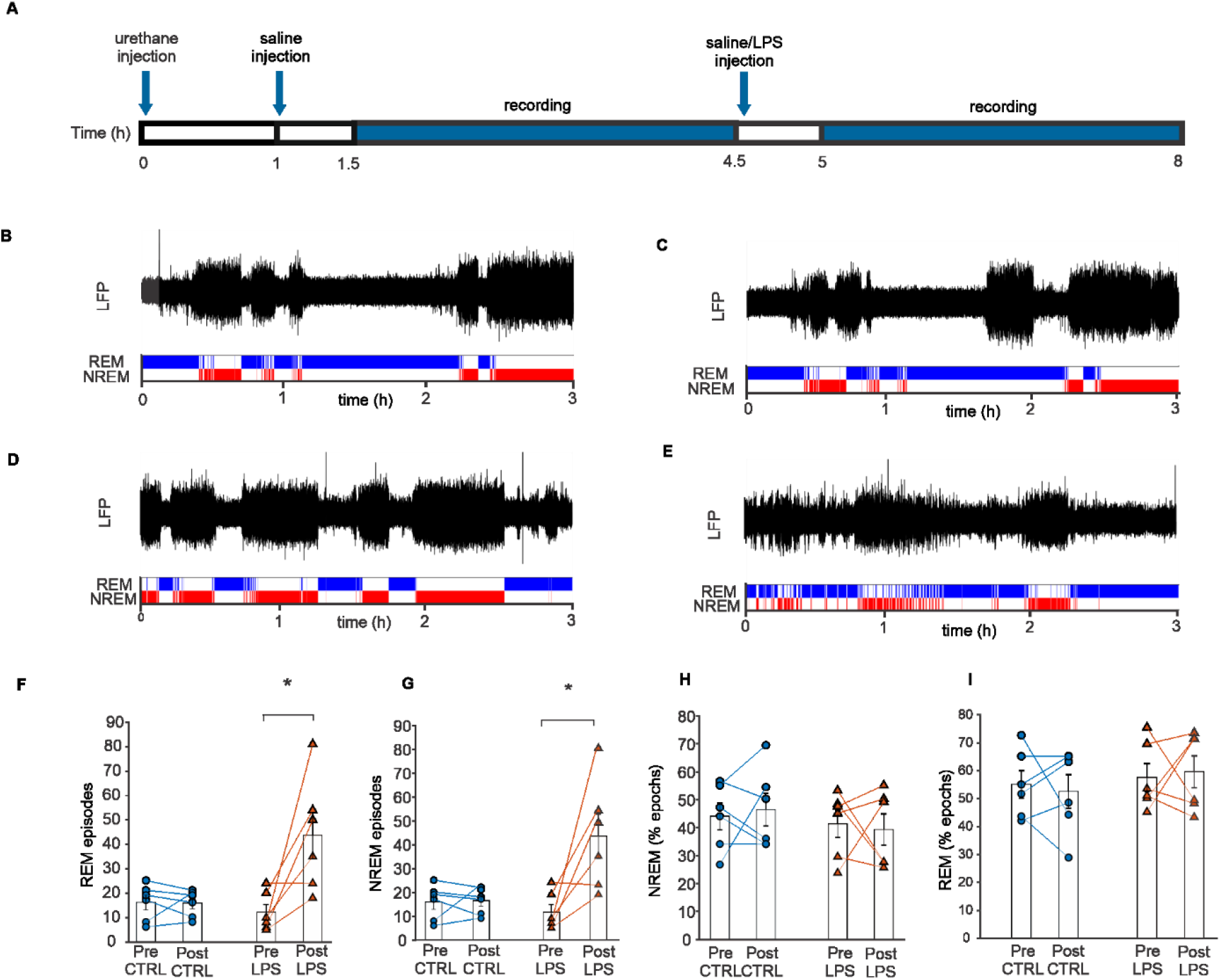
LPS causes state fragmentation in rats without affecting the total time spent in either state. **A.** Experimental design. Rats in urethane anesthesia were injected with saline and baseline LFP was recorded for 3-hours, followed by a single dose of LPS (LPS, n=6) or saline (CTRL, n=6) and another a 3-hour LFP recording. **B-E.** Representative traces and hypnograms showing the REM-like state in blue, NREM in red. Sleep states were long and stable before (B) or after saline injection (C) in CTRL group and before the LPS injection (D) in the LPS group. After LPS injection (E), sleep states were heavily fragmented. F. Number of REM episodes before and after treatment. Each data point shows the number of REM episodes in individual CTRL and LPS animals pre and post injection (CTRL, n = 6; LPS, n = 6). Bars show mean ± s.e.m. **G.** Number of NREM episodes in CTRL and LPS animals pre and post injection. **H.** Total time spent in REM as percentage of recording time in CTRL and LPS animals pre and post injection. **I.** Total time spent in NREM as percentage of recording time in CTRL and LPS animals pre and post injection. *p < 0.05.

### 2.3. Electrode implantation surgery

Local field potentials were recorded from the rats implanted with eight independently movable tetrodes in the CA3 region of the hippocampus. Each tetrode consisted of four twisted 17-μm polyimide-coated platinum-iridium wires coated with platinum in order to reduce the impedance to 120 −200 kΩ at 1 kHz. Rats were anaesthetized using a mixture of ketamine (Narkamon, 100 mg/kg, i.p.) and xylazine (Rometar, 10 mg/kg, i.p.) and 1.5-2% isoflurane in O_2_. They were fixed in a stereotaxic frame and body temperature was maintained at 37 °C using a heating pad. An incision was made in the scalp to expose the skull, after which a craniotomy was made over the dorsal hippocampus (3.8 mm AP, 3.2 mm lateral of bregma). After removal of the dura mater, the tetrode bundle was carefully lowered into the cortex. Individual tetrodes were slowly lowered into CA3 over the course of a 2-week recovery period. The hyperdrive was fixed to the skull using dental acrylic and stainless-steel screws. One screw, located above the frontal cortex, served as a reference. Rats were given carprofen (Rimadyl, 5 mg/kg, s.c.) and Marbofloxacin (Marbocyl, 5 mg/kg, s.c.) during recovery.

### 2.4. Recording procedure

Rats used were previously involved in behavioral experiments for 2-3 weeks after surgery. For the experiments described in this paper, rats were anesthetized for recording using urethane (1.5 g/kg, i.p., Sigma). An hour after urethane injection, hippocampal local field potentials were amplified using an Intan RHD2132 headstage amplifier, digitized and recorded using an OpenEphys recording system at a sampling rate of 2000 Hz (Siegle et al., 2017).

### 2.5. Data processing and analysis

#### 2.5.1. Signal processing

For vigilance state classification and later state-space analysis, a sliding window FFT analysis (2 s window, 1 s step) was performed on separately for all recorded channels using Welch’s method in MATLAB 2014b (Hamming window, 50% overlap, 0.25 Hz resolution), yielding 1 epoch per second. Then, spectral ratios were calculated for overall power in two overlapping frequency ranges for each time window. Ratio 1 was calculated as R1 = (1-2 Hz)/(1-9 Hz) and Ratio 2 was calculated as R2 = (1-15 Hz)/(1-45 Hz). Frequency ranges were chosen based on literature and on the approximate frequencies of delta and theta peaks in our recorded baseline spectra (Diniz Behn et al., 2010; Gervasoni et al., 2004). Normalized signal amplitude was calculated using the same sliding window approach, where mean absolute signal amplitude was calculated for each window and normalized to mean absolute signal amplitude for the entire recording. The resulting power and amplitude time series for all channels were then combined into a single time series per rat for each of these variables using PCA. The first principal component was used for further analyses. Epochs that contained artefacts were excluded from further analysis (0.18 ± 0.06% of recording time). State labels for these epochs were set to be identical to the preceding, artefact-free epoch prior to smoothing.

#### 2.5.2. Vigilance state classification and episode detection

Brain activity under urethane anaesthesia showed two distinct alternating states: a NREM-like state that was dominated by low frequency, high-amplitude waves, and a REM-like state with faster activity and a lower signal amplitude. Epochs were automatically classified as belonging to one of these states based on the calculated principal component values for R1, R2, and amplitude using k-means clustering. Inclusion of the amplitude parameter in the clustering procedure ensured reliable state identification, even when spectral ratios were affected by experimental treatment.

After initial clustering, ultra-short periods of NREM-like or REM-like activity were removed: a state transition was only considered if the first epoch of the new state was followed by at least 3 more epochs of the same state. Otherwise, the epoch was labelled as belonging to the preceding state. This smoothing procedure ensured that any unrealistically short, artefactual state changes caused by normal within-state signal variability were removed.

NREM-like and REM-like episodes were calculated as periods of each state that were at least 10 epochs long, and were followed by at least 10 epochs of the othe*r* state. Short episodes were considered as 20 – 120 s long, whereas episodes of 600 s or more were considered long. Epoch-to-epoch transition probability was calculated based on the percentage of epoch that was followed by the same or a different state. Episode transitions were characterized by fitting a logistic curve using MATLAB’s fit function with the following equation f(x) = offset + (range / (1 + e^−slope^*^x^)). Curves were fit to a 30-s period of LFP trace centered on the episode state transition after smoothing using a 5-s window moving average.

#### 2.5.3. State-space analysis

Inflammation-related spectral changes were analysed in a 2-dimensional state-space based on R1 and R2. This type of analysis may reveal within- and between state dynamics that are not captured using less sensitive single-band approaches (Diniz Behn et al., 2010; Gervasoni et al., 2004). Cluster positions were defined using the median pc1R1 and pc1R2 values for each state.

Within-state stability was analyzed using state-space velocity. Velocity was defined as the Euclidian distance between two subsequent epochs. As such, overall velocity was calculated as v = √((pc1R1_n+1_-pc1R1_n_)^2^+(pc1R2n+1-pc1R2_n_)^2^), and velocity along a single dimension such as R1 simply as v_R1_ = pc1R1_n+1_ – pc1R1_n_

#### 2.5.4. Analysis of periodic and aperiodic spectrum components

To further investigate how power spectrum changes lead to the observed state-space effects, aperiodic and periodic components of the power spectrum were parametrized using the FOOOF algorithm (v. 1.0.0) in Python 3.7 (Donoghue et al., 2020; Haller et al., 2018). First, average power spectra for each state were calculated from the sliding window FFT described earlier. One representative channel was analysed per rat and the same channel was used for the pre- and post-injection time points. For each spectrum, following aperiodic fit AP(f) = 10^b^ * (1/(k+f×)) fit (AP), where f is frequency, b is offset, k is the knee parameter, and χ is the spectrum slope. The knee parameter represents the bending point where the aperiodic fit transitions from horizontal to negatively sloped. Knee frequency is dependent on the value of k and spectrum slope χ and was calculated as k_freq_ = k^(1/ χ)^. Periodic components of the spectrum, representing putative oscillations, were modelled as Gaussian curves over and above the aperiodic background spectrum. These oscillations each have a center frequency (c), peak width (w), and center peak height (a), yielding the following for each oscillation frequency f G(f) = a * exp(-(f-c)^2^/(2*w^2^)).

### 2.6. Cytokine quantification and histology

After recording, blood samples were collected and serum was separated by centrifugation at 1,000 × g for 10 min. Levels of IL-1β were quantified using ELISA kit according to manufacturer’s protocol the frequency range from 1 to 45 Hz was used with the following algorithm settings: peak width limits: 0.5 and 12, maximum number of peaks: 6, minimum peak height: 0.2, peak threshold: 2.0, and aperiodic mode: knee. Aperiodic spectral components were modelled using the (RAB0277, Sigma).

After this, rats were killed using an overdose of sodium pentobarbital (50 mg/kg) and perfused transcardially with ringer solution followed by 4% paraformaldehyde in phosphate-buffered saline. Brains were collected and cut in 50μm thick coronal sections. Electrode placement was verified using Nissl staining. Electrode traces from electrodes outside of hippocampal CA3 were excluded from analysis.

## 3. Results

### 3.1. LPS increases serum IL-1β

To confirm the presence of systemic inflammation in LPS-injected animals, IL-1β levels were measured using ELISA. Serum IL-1β concentrations in LPS-injected rats were at least 3 times higher than in saline-injected controls. (398 ± 32.24 pg/ml vs 122 ± 63.64 pg/ml, n = 6, independent sample t-test t(10) = 3.88, p < 0.01, Fig. S1).

### 3.2. LPS injection causes altered sleep state structure

Under urethane anaesthesia rats showed two distinct patterns of brain activity: a NREM-like state that was dominated by high amplitude slow waves, and a REM-like state with low-amplitude, higher frequency activity (Fig. 1B-E). The states were long and stable at baseline and in saline-injected controls (Fig. 1B-D), but much shorter after LPS injection (Fig. 1E). The total number of episodes per recording period was significantly affected by LPS injection (rmANOVA; group effect: F(1, 10) = 5.14, p = 0.04; time effect F(1, 10) = 8.64, p = 0.01; group*time interaction: F(1, 10) = 8.55, p = 0.01, Fig. 1F-G). Post-hoc analysis showed a significantly higher number of episodes after LPS injection (87 ± 20 episodes) compared to before injection (24 ±16 episodes). As a result, episode duration was also significantly altered (rmANOVA; group effect F(1, 10) = 0.03, p = 0.85; time effect: F(1, 10) = 10.46, p < 0.01; group*time interaction: F(1, 10) = 5.58, p = 0.04). Post-hoc analysis showed that episodes in the LPS group were significantly shorter after LPS injection (167 ± 36 s) than at baseline (694 ± 153 s). However, despite overall changes in episodes number, the time spent in NREM (rmANOVA; time effect: F(1, 10) = 0.002, p = 0.96; group effect: F(1, 10) = 0.77, p = 0.39; group*time interaction : F(1, 10) = 0.199, p = 0.66) and in REM (rmANOVA; time effect: F(1, 10) = 0.002, p = 0.96, group effect: F(1, 10) = 0.77, p = 0.39; group*time interaction: F(1, 10) = 0.199, p = 0.66) was not significantly changed by LPS injection (Fig. 1H-I).

Extensive sleep instability after LPS injection was further apparent in episode length distribution (Fig. 2A). At baseline and after saline injection rats had few short episodes (20s −120s) and a relatively high number of long episodes (≥600 s). However, after LPS injection, brain activity patterns consisted of many short episodes and only a few long ones. These effects were present in both NREM and REM.

**Fig. 2.**
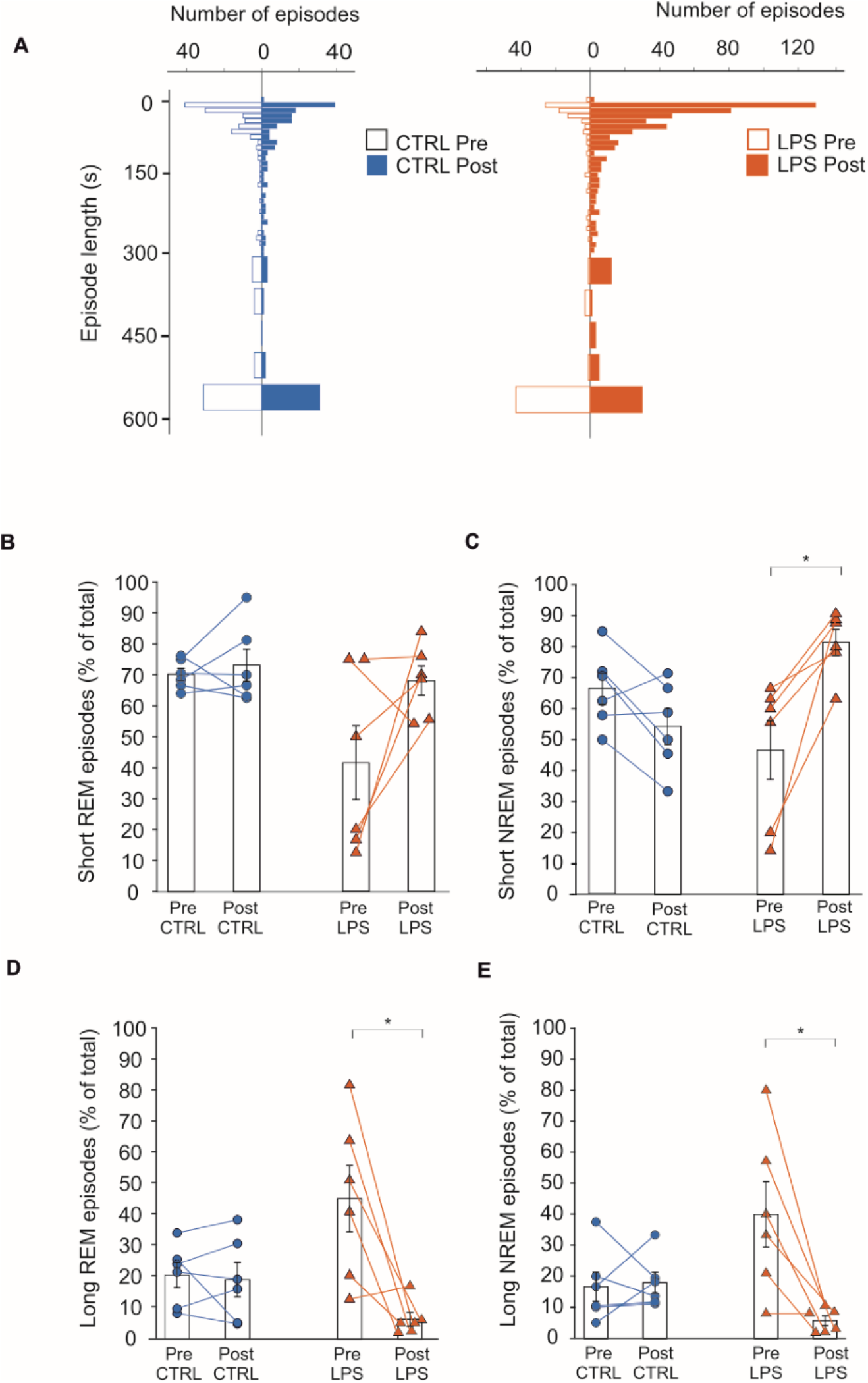
LPS leads to an increase in short and to a decrease in long sleep state episodes. A. Histograms depicting the overall distribution of episode lengths in CTRL (left) and LPS rats (right), pre and post injection. Each bar in the histogram represents the number of episodes per 10- or 50-s bin. B-C. Proportions of short (20s-120s) episodes of REM (B) or NREM (C) in CTRL and LPS rats, pre and post injection (CTRL, n = 6; LPS, n = 6) in the recording. D-E. Proportions of long (≥600s) episodes of REM (D) or NREM (E) in CTRL and LPS rats, pre and post injection. Bar graphs show mean ± s.e.m. *p < 0.05.

The relative number of short episodes in the recording period showed a significant time effect (rmANOVA; F(1, 10) = 4.75, p = 0.05) and group*time interaction (F(1, 10) = 8.63, p = 0.01), but no significant group effect (F(1, 10) = 1.08, p = 0.32). Post-hoc analysis showed an increase in the short episodes in the LPS group post injection (75 ± 4.6% of episodes) compared to pre-injection (44.1 ± 10.9% of episodes).

When quantified separately, relative amounts of short episodes in NREM were significantly higher in NREM after LPS injection, but not in REM (Fig. 2B, C). Short NREM episodes showed a significant time effect (rmANOVA; F(1, 10) = 4.67, p = 0.05) and a non-significant group effect (F(1, 10) = 0.25, p = 0.62). There was a significant effect of group*time interaction (F(1, 10) = 19.78, p <0.01). Post-hoc analysis showed a significant increase in the percentage of short NREM episodes in the LPS group post injection (81.44 ± 4.58% of NREM episodes vs 46.59 ± 10.37% pre-injection). In REM, there was a significant group effect (rmANOVA; F(1, 10) = 6.58, p = 0.02) but a non-significant time effect (F(1, 10) = 4.01, p = 0.07) and group*time interaction (F(1, 10) = 2.56, p = 0.14). Post-hoc analysis showed a significant baseline difference between controls (70.12 ± 1.94%) and the LPS group (41.52 ± 11.88%).

The percentage of long episodes in the recording period was also affected by LPS injection, but showed opposite effects to those observed for short episodes. The total percentage of all long episodes showed a significant effect of time (F(1, 10) =9.39, p=0.01) and group*time interaction (F(1, 10) = 9.18, p = 0.01), but no significant group effect (F(1, 10) = 1.00, p = 0.34). Post-hoc analysis showed a significant decrease in the percentage of long episodes in the LPS group post-injection (6.07± 1.55% of episodes) compared to pre-injection (42.20 ± 10.43% of episodes), although absolute numbers of long episodes are quite low.

Effects of LPS injection on NREM and REM states separately paralleled those found in the overall episode length distribution (Fig. 2D,E). The percentage of long NREM episodes showed a significant effect of time (rmANOVA; F(1, 10) = 8.28, p = 0.01) and group*time interaction (F(1, 10) = 9.40, p = 0.01). There was no significant group effect (F(1, 10) = 0.79, p = 0.39). Post-hoc analysis showed a decrease in the percentage of long NREM episodes after LPS injection (5.90 ± 1.55% of NREM episodes) compared to baseline (39.97 ± 10.48% of NREM episodes). Long REM episodes also showed a significant time effect (rmANOVA; F(1, 10) =9.78, p=0.01) and group*time interaction (F(1, 10) = 8.39, p = 0.01), but no significant group effect (F(1, 10) = 0.89, p = 0.36). Post-hoc analysis showed a significantly lower percentage of long REM episodes in the LPS group post-injection (6.51 ± 2.12% of REM episodes) compared to pre-injection (44.16 ± 10.42% of REM episodes, Fig. 2D,E). These observed effects on episode number and duration point to instability in both the REM- and NREM-state.

State instability was also seen on the epoch-to-epoch level, where the probability of a state transition from one epoch to the next was similarly increased for NREM to REM transitions (rmANOVA; time effect: F(1, 10) = 16.72, p <0.01; group effect: F(1, 10) = 4.61, p = 0.06; group*time interaction: F(1, 10) = 10.53, p<0.01) and REM to NREM transitions (rmANOVA; time effect: F(1, 10) = 16.12, p<0.01; group effect: F(1, 10) = 4.77, p = 0.05; group*time interaction: F(1, 10) = 10.68, p<0.01). Post-hoc analysis showed a significantly higher probability of state transition from NREM to REM (0.17 ± 0.05% pre vs. 0.62 ± 0.10% post) and REM to NREM (0.17 ± 0.05% pre vs. 0.62 ± 0.09% post) in the LPS group post injection (Fig. S2). State transition probabilities remained at baseline levels in the CTRL group (NREM to REM: 0.22 ± 0.05% vs 0.27 ± 0.05%; REM to NREM: 0.27 ± 0.05% vs 0.27 ± 0.05%)

Although a higher number of state transitions was found after LPS injection, the characteristics of these transitions were not significantly different from those in controls or at baseline (Fig. S3). Thus, LPS injection leads to instability of NREM and REM to a similar degree, without affecting time spent in either state, or the characteristics of transitions between the states.

### 3.3. LPS leads to increased spectral similarity between REM and NREM

Changes in spectral characteristics were assessed based on two spectral power ratios: Ratio1 (R1, 1-9 Hz/1-2 Hz) and Ratio 2 (R2, 1-45 Hz/1-15 Hz), which are distinct in the two observed sleep-like states (Fig. 3). State-space analysis of REM and NREM based on these spectral power ratios resulted in two distinct state clusters of epochs in the control group and LPS group at baseline and after injection (Fig. 4A-D).

**Fig. 3.**
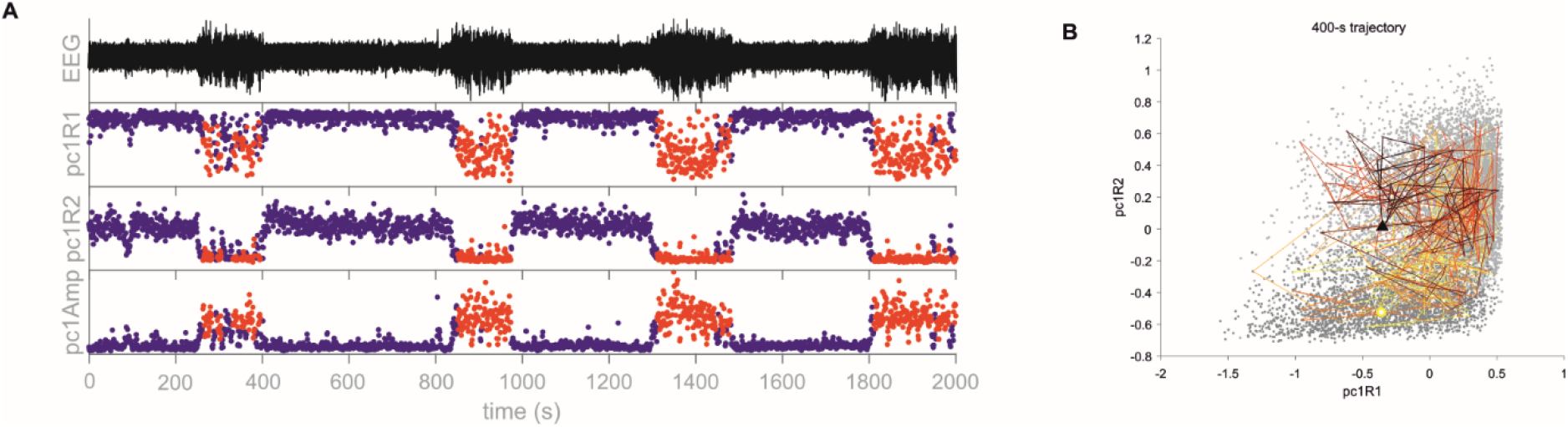
NREM and REM states are distinct in spectral state space. **A.** First principal component of spectral ratios R1 and R2 in time, as well as the amplitude component used for state classification, with the accompanying EEG trace. Epochs classified as NREM are shown in red, epoch classified as REM are shown in blue. **B.** Example of a 400-s long trajectory in 2-D state space. The shown trace starts in the NREM cluster (circle) and eventually ends in the REM-like cluster after covering much of the recording’s state-space. Epochs that are not part of this trajectory are shown in grey.

**Fig. 4.**
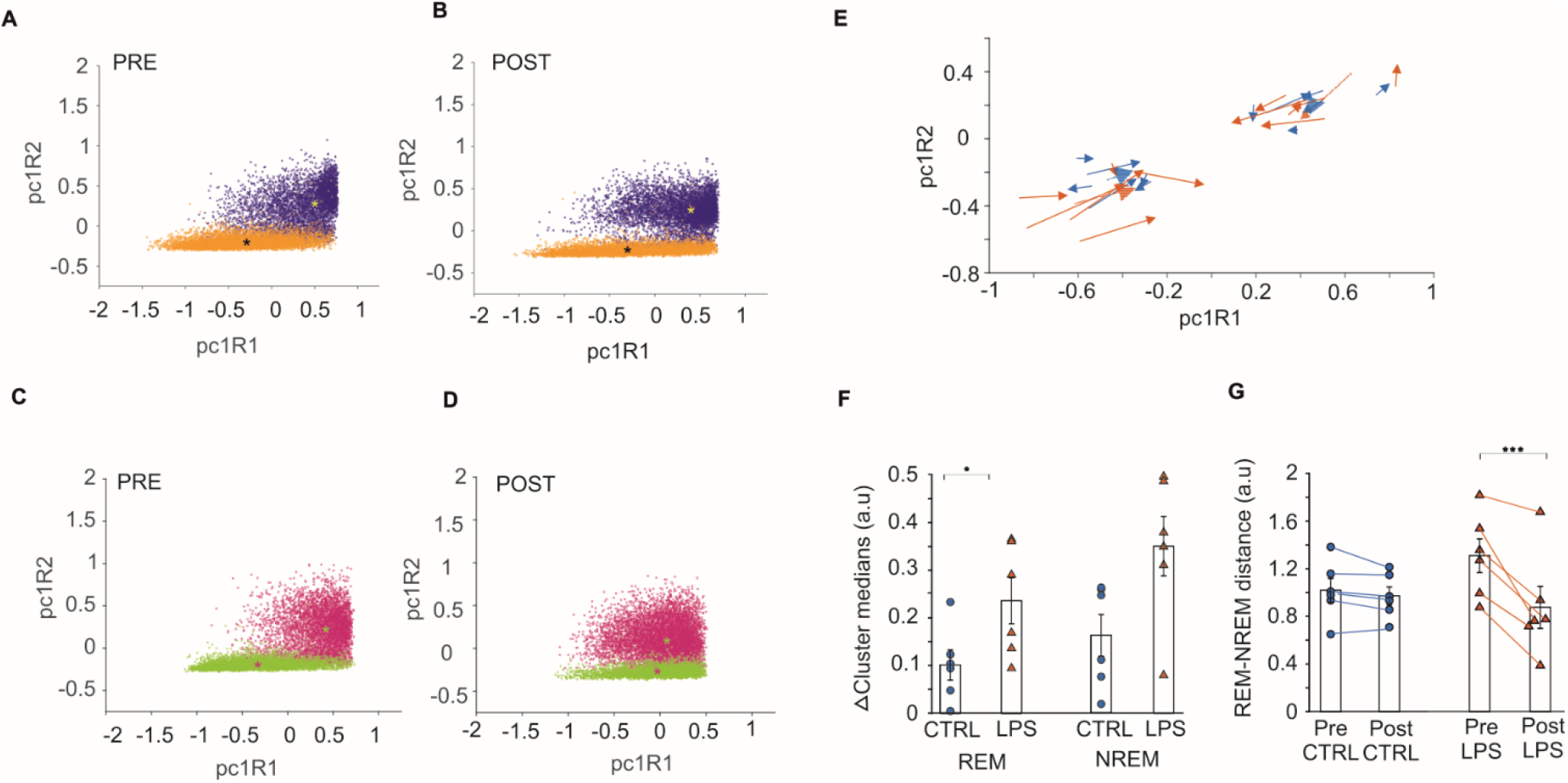
LPS leads to reduced cluster distance between REM and NREM states. **A-D**. Scatter plots showing REM and NREM epochs in 2-D state space in CTRL (A, B) and LPS (C, D) groups, pre and post injection. The asterisk marks the cluster median. **E.** Quiver plot showing the directional shift of the REM and NREM cluster medians before and after injection in CTRL (blue) and LPS animals (orange). The starting point of each arrow shows cluster medians before the injection, and the arrow head the medians after injection. **F**. Size of the shift in cluster medians in REM and NREM. Each data point represents change in cluster position after the treatment in individual CTRL or LPS animals. Bars show mean ± s.e.m. **G.** The distance between REM and NREM clusters in CTRL or LPS animals pre and post injection. *p<0.05; ***p<0.001.

After LPS injection (Fig. 4D), the location of the NREM and REM clusters within state-space shifted, resulting in a decreased inter-cluster distance. The magnitude and direction of the changes in cluster location after LPS injection and in controls is shown in Fig. 4E.

The centroid of REM cluster shifted significantly in LPS group (0.23 ± 0.05 a.u.) compared to control group (0.10 ± 0.03 a.u., t(10) = 2.33, p = 0.04, Fig. 4F). The direction of mean resultant vector is towards the origin for REM cluster (Fig. 4E). In case of NREM cluster, there was no significant difference between the cluster centroids of control (0.16 ± 0.04 a.u.) and LPS group (0.35 ± 0.06 a.u., t(10) = 1.45, p = 0.17). The mean resultant vector however is directed away from origin, towards the REM cluster (Fig.4E).

As a result of the observed shifts in cluster medians, the total distance between the two clusters decreased (Fig. 4G). We calculated the total distance between REM and NREM clusters before and after injection in control and LPS groups and observed a significant time effect (F(1,10) = 26.47, p < 0.001) and group*time interaction (F(1,10) =17.09, p <0.01). There was no group effect (F(1, 10) = 0.3, p = 0.59). Post-hoc analysis showed a significantly smaller distance between REM and NREM clusters in the LPS group post-injection (0.87± 0.19 a.u.) compared to baseline (1.30 ± 0.15 a.u.). The decreased inter-state cluster distance indicates increased spectral similarity between the states.

### 3.4 State similarity in low spectrum frequencies

The decreased distance between REM and NREM clusters could be the result of changes in R1, R2, or in both. By decomposing the total distance inter-cluster distance into R1 and R2 components, we determined which spectral range most affected post-LPS brain activity. Distance between the REM and NREM clusters was significantly decreased along R1 (rmANOVA; time effect F(1, 10) = 39.26, p < 0.001, group effect F(1, 10) = 0.03, p = 0.84, group*time interaction, F(1, 10) = 27.55, p < 0.001). Post-hoc analysis showed that the distance between clusters was significantly smaller in the LPS group post injection (0.69 ± 0.20 a.u.) compared to baseline (1.13 ± 0.14 a.u., Fig. S4A). By contrast, there was no significant change in the distance between the clusters along R2 (rmANOVA; time effect, F(1, 10) = 2.85, p = 0.12, group*time interaction, F(1, 10) = 1.59, p = 0.23, group effect, F(1, 10) = 1.64, p = 0.22, Fig. S4B).

Hence, LPS-mediated spectral similarity between REM and NREM is mostly the result of changes in the 1 to 9 Hz frequency range.

### 3.5. NREM contributes more to spectral similarity in lower frequencies than REM

After observing a reduced distance between REM and NREM clusters along R1, we investigated if this shift was state-specific or if both states contributed. In REM, we observed no significant changes in median R1 values (rmANOVA; time effect: F(1, 10) = 3.16, p = 0.10, group effect: F(1, 10) = 0.0002, p = 0.98, time*group interaction: F(1, 10) = 4.31, p = 0.06, Fig. S4C).

In NREM however, we found a significant time effect (F(1, 10) = 15.02, p <0.01) and time*group interaction (F(1, 10) = 7.27, p = 0.02). There was no significant group effect (F(1, 10) = 0.13, p = 0.72). Post-hoc analysis showed a significant increase in R1 medians in the LPS group post-injection (0.61 ± 0.09 a.u.) compared to baseline (0.34 ± 0.09 a.u., Fig. S4D).

### 3.6. LPS causes increased within-state instability in REM and NREM

The observed state fragmentation and altered REM-NREM dynamics could be caused by inflammation-related state instability. Here, we used distance between subsequent epochs in state-space, or within-state velocity, as a measure of stability.

Trajectories of short time-series sampled from the EEG signal that start or end near a cluster median generally remain within the limits of the surrounding (Fig. S6), but show high second-to-second variability. Epochs that are neighbouring in time are not necessarily located in close proximity within the spectrum-based state space. This is the case in both the LPS and control groups, although longer trajectories to and trajectories from the other state after LPS injection. This is in part due to an increased number of state transitions that may be captured in the longer timeframe, where a trajectory starting in one state ends up within the other. Mostly, however, the increased overlap is due a combination of decreased inter-cluster distances and the observed spatial variability, where longer trajectories end up covering much of a state cluster even without leaving it.

Velocities along R1 were not significantly affected by LPS injection in either REM (rmANOVA; time effect: F(1,10) = 1.20, p = 0.29; group effect: F(1,10) = 0.80, p = 0.39; time*group interaction: F(1,10) = 0.01, p = 0.92, Fig. S5A), or NREM (time effect: F(1,10) = 0.92, p = 0.36; group effect: F(1,10) = 0.86, p = 0.37; time*group interaction: F(1,10) = 0.0005, p = 0.98, Fig. S5B).

However, velocities along R2 were significantly increased in both REM and NREM after LPS injection. In REM, we observed a significant time*group interaction (F(1,10) = 7.93, p = 0.01) but no significant effect of group (F(1,10) = 0.96, p = 0.34) or time (F(1,10) = 1.17, p = 0.30) (Fig. S5C). Post-hoc analysis showed significantly higher velocities in the LPS group after injection compared to baseline (0.087 ± 0.03 a.u. vs. 0.074 ± 0.03 a.u.). The effects on R2 velocities in NREM were similar: there was a significant time*group interaction (F(1,10) = 7.15, p = 0.02), but no significant effect of group (F(1,10) = 0.96, p = 0.34) or time (F(1,10) = 1.17, p = 0.30, Fig. S5D). Post-hoc analysis showed no significant differences.

### 3.7 Effects of LPS on periodic and aperiodic power spectrum components

To investigate possible sources of the observed changes in R1, power spectra were analysed for representative channels. NREM spectra showed a marked reduction power below 3 Hz in LPS-injected rats, but not in controls (Fig. 5A). REM spectra showed the opposite effect: increased power in the < 3 Hz range, as well as a smaller increase in the 7-9 Hz range (Fig. 5B). Particularly in REM, these changes were quite variable. Power spectra of rats in the control group remained largely stable and at baseline levels.

**Fig. 5.**
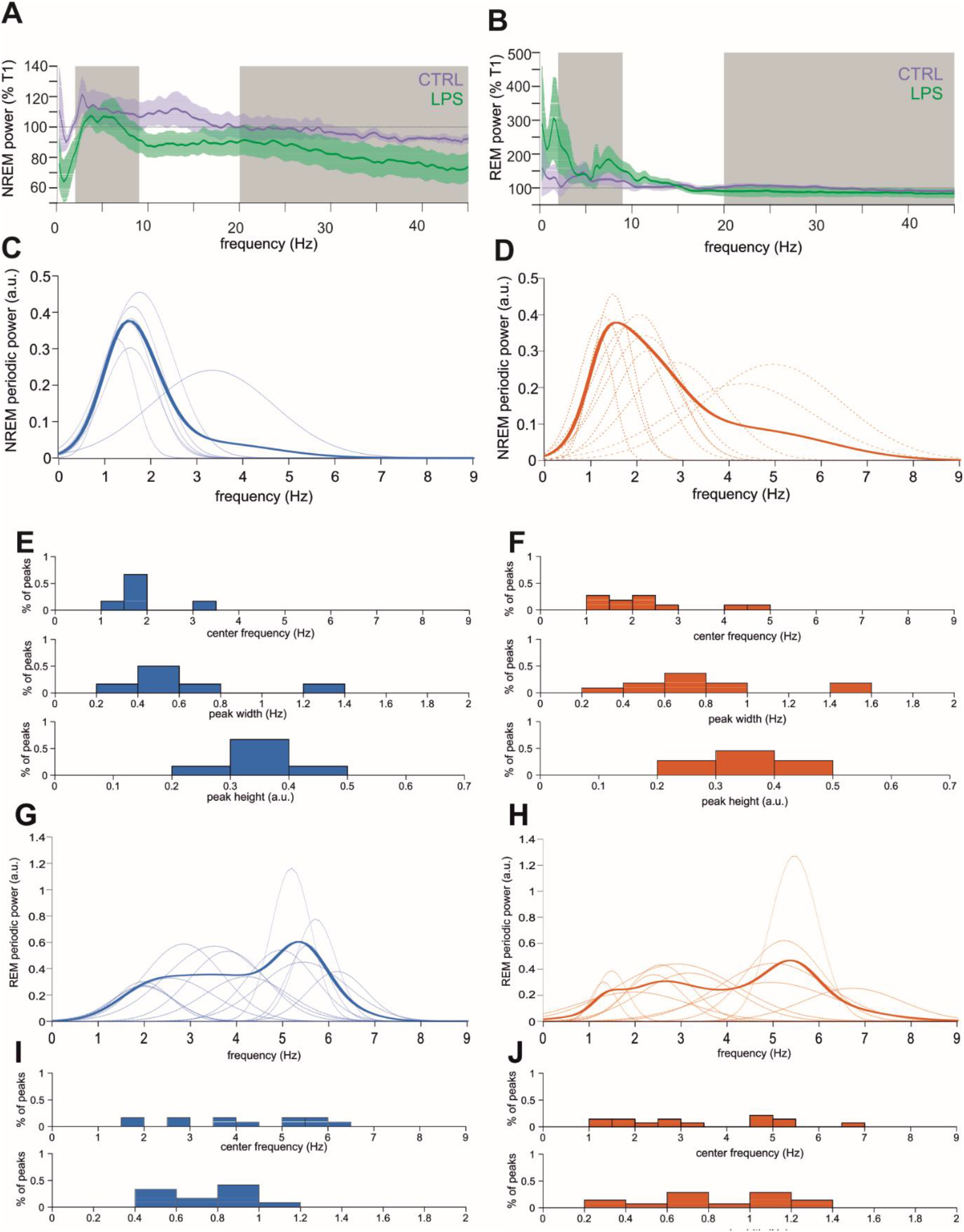
LPS-related changes in NREM and REM power spectra. **A.** NREM power spectra of control (purple) and LPS-injected (green) rats as percentage of pre-injection baseline (T1). Grey areas indicate limits of the spectral ratios used for state-space analysis. **B.** REM power spectra of control (purple) and LPS-injected (green) rats as percentage of pre-injection baseline (T1). Grey areas as in (A). from either state show more overlap with **C-D.** NREM oscillations in the 1-9 Hz frequency range prior (C) and after (D) the LPS injection. Dotted lines show individual values solid line shows the mean oscillatory power. **E-F.** Distribution of NREM oscillation characteristics before (E) and after (F) the LPS injection. **G-H.** REM oscillations in the 1-9 Hz frequency range prior (G) and after (H) the LPS injection. Lines as in (C). **I-J.** Distribution of REM oscillation characteristics before (I) and after (J) the LPS injection.

The observed changes in spectral power ratios and spectral power distribution could be the result of alterations in EEG oscillations and in the background (aperiodic) components of the power spectrum. To better understand the observed changes in R1 in REM and NREM after LPS injection, aperiodic and periodic components of the power spectrum for each state were modelled using FOOOF in the LPS group. Overall model fits were good for NREM spectra (R^2^ = 0.995 ± 0.002, fit error = 0.031 ± 0.006) and REM spectra (R^2^ = 0.987 ± 0.008, fit error = 0.035 ± 0.007). Model fitting was not negatively affected by LPS injection (post-LPS fits in NREM: R^2^ = 0.996 ± 0.002, fit error = 0.029 ± 0.008, and REM: R^2^ = 0.985 ± 0.011, fit error = 0.035 ± 0.006).

Similar to non-anaesthetized recordings (Leemburg et al., 2018), REM spectra had overall slightly shallower slopes than the NREM spectra (pre-LPS REM vs. NREM exponents: 1.42 ± 0.28 vs. 1.95 ± 0.18; Fig. S7A-B). This was also the case after LPS injection (post-LPS REM vs. NREM exponents: 1.67 ± 0.22 vs. 2.09 ± 0.19; Fig. S7A-B)

The aperiodic components of the NREM spectra were not significantly affected by LPS injection (Fig. S7). Spectrum slopes remained stable at 107.5 ± 3.3% of pre-LPS slope values (Wilcoxon signed rank test, W = 21, p = 0.09 after Bonferroni correction, Fig. S7C), as did model offsets (102.15 ± 0.49%, W = 21, p = 0.09 after Bonferroni correction). Knee frequencies were more variable than the other parameters (pre-LPS: 1.83 ± 0.38 Hz, post-LPS: 2.68 ± 0.61 Hz) and showed a slight increase after LPS injection, but this effect was not significant (144.39 ± 11.36%, W = 21, p = 0.09 after Bonferroni correction, Fig. S7C).

Effects of LPS on aperiodic components in REM were similar to those observed in NREM (Fig. S7B). Like in the NREM spectra, REM slopes showed no significant changes after LPS injection (127.39 ± 15.09% of pre-LPS, W = 16, p = 0.94 after Bonferroni correction, Fig. S7D). NREM knee frequencies (pre-LPS: 5.37 ± 2.68 Hz, post-LPS: 5.49 ± 2.00 Hz, W = 15, p = 1.31 after Bonferroni correction, Fig. S7D) and spectrum offsets were likewise not significantly affected (pre-LPS: 4.13 ± 3.44 a.u., post-LPS: 4.42 ± 0.35 a.u., W = 16, p = 0.94 after Bonferroni correction, Fig. S7D). They remained at 170.83 ± 77.38% and 107.63 ± 5.45% of baseline, respectively.

In pre-LPS recordings, NREM spectra showed one major periodic component: an oscillation in the delta frequency range with a mean center frequency of 1.56 ± 0.06 Hz and peak widths between 0.2 and 0.8 Hz (Fig. 5C-D). After LPS injection, center frequencies of this oscillation increased and peak widths became more variable (Fig. 5E-F). Overall, the main oscillatory component in NREM became faster and more variable after LPS injection. As such, the amount of spectral power in frequencies higher than 2 Hz increased, resulting in the observed shift in Ratio 1 in NREM.

REM spectra showed multiple oscillations: a delta-like oscillation with a peak frequency in the 1 - 2 Hz range, similar to the one found in NREM, and a second theta-like oscillation with a peak frequency around 5 – 6 Hz (Fig. 5G-H). These oscillations remained present after LPS injection, but more oscillations were found to have slower peak frequencies, and more oscillations had peak frequencies lower than 2 Hz. Additionally, REM peak widths became more variable, and overall peak heights tended to be slightly lower (Fig. 5I-J). This slowing of the oscillatory components in REM resulted in relatively more power in the 1-2 Hz range, while the power in frequencies over 2 Hz was decreased. This effect, opposite in direction to that found in the NREM spectrum, resulted in the decreased spectral distance between NREM and REM clusters observed in state-space.

## 4. Discussion

In the present study, we used the lipopolysaccharide model of sepsis in rats to characterize effects of severe generalized inflammation on brain oscillation state dynamics. We found profound alteration of EEG state dynamics in LPS-injected animals, mainly expressed as instability between the NREM-like and REM-like states: the states alternated considerably more frequently as the episode lengths were shortened by approximately 75%. Additionally, the time spent in either state was not affected by LPS in our study, indicating that the observed effect is unlikely to be the result of selective suppression of one of the states. Rather, it indicates that increased switching between the slow-wave dominated NREM-like state and the higher frequency, low amplitude REM-like state was a result of the generalized inflammatory response. In the bistable state (NREM-like, REM-like) system under urethane, this resembles the fragmentation of NREM by wakefulness rather than by REM sleep in the non-anaesthetized system and might provide thus a complexity-reduced model for the sleep fragmentation.

Sleep disturbances are a frequent complication of acute systemic inflammation in early- and developed-stage sepsis patients of largely unknown pathophysiology, and may persist in post-sepsis syndrome (Richards and Bairnsfather, 1988; Weinhouse et al., 2009; Weinhouse and Schwab, 2006). In the acute stage, state fragmentation presents as many short sleep episodes that are widely distributed across the 24-hour period (Baracchi et al., 2011), indicating possible disturbances in both the homeostatic and circadian sleep regulatory systems: changes of central melatonin release rhythms (Bourne et al., 2008), as well as circadian regulation of suprachiasmatic and HPA-axis activity (Beynon and Coogan, 2010; Carlson et al., 2006; Cavadini et al., 2007). Although the main molecular drivers of sleep homeostasis are not yet known, inflammation has been shown to affect expression of BDNF and adenosine 2A receptors (Cui et al., 2021; Ohta and Sitkovsky, 2001; Pereira de Souza Goldim et al., 2020), both of which have been linked to sleep regulation (Lanza et al., 2022, p. 4; Reichert et al., n.d.; Schmitt et al., 2016). Additionally, cytokine signaling plays an important role in regulating sleep pressure under normal circumstances (Opp, 2005).

To further investigate possible causes of oscillatory state switching within the hippocampal network we used a power spectrum-based state-space approach. By plotting each epoch according to two spectral ratios and classifying as NREM or REM based on their position within the resulting state-space, we were able to study the relation between the states and the within-state dynamics. Similar state-space approaches have previously been used to study vigilance state dynamics in humans, mice, and rats (Diniz Behn et al., 2010; Gervasoni et al., 2004; Imbach et al., 2016; Schoch et al., 2017). The distance between NREM and REM clusters was significantly reduced in septic animals. Such increased similarity between distinct oscillatory states has not been described in sepsis patients or animal models so far. Similar decreases in inter-state distances have been observed in narcolepsy patients and in orexin knockout mice, expressing a narcolepsy-like phenotype that is characterized by a high level of state fragmentation (Diniz Behn et al., 2010; Schoch et al., 2017). Reduced inter-state distances in state-space could therefore be a general characteristic of high state instability, and may indicate altered attractor dynamics between stable states, facilitating the state transitions.

We found that decreased state-space distance between NREM and REM clusters was driven by opposing effects of LPS on these states within the lower frequency ratio R1. In NREM, this consisted of decreased delta power, as well as a shift towards faster oscillations within that frequency range. Decreases in NREM delta power have previously been described in sepsis in rats (Baracchi et al., 2011), although effects of lower doses of LPS on NREM delta power are less consistent (Kapás et al., 1998; Lancel et al., 1995). Increased delta and theta power, with high inter-individual variability, as well as a general slowing of oscillatory activity in the REM-like state under urethane is in line with slowing of EEG activity during awake, or wake-like states, described in sepsis patients (Götz et al., 2016; Rosengarten et al., 2012; Straver et al., 1998; Urdanibia-Centelles et al., 2021). Increased slowing was associated with poor outcomes, including delirium (Urdanibia-Centelles et al., 2021) and death (Yamanashi et al., 2021). In animal models, increased wake delta power (Kafa et al., 2010), slowed EEG (Kurita et al., 2010), and reduced theta frequencies have been described (Mamad et al., 2018). Slowed EEG may persist in former sepsis patients (Semmler et al., 2008), but long term effects in animal models are less consistent (Rong Gao et al., 2017; Ji et al., 2020; Wang et al., 2020). It is not clear how much the previously described power spectrum changes are due to altered background spectrum or specific oscillatory activity, as the peak frequencies within spectral power bands were typically not analyzed. The occurrence of isolated delta and theta waves in the wake EEG of septic patients suggests that they are at least in part related to oscillations (Bolton, 1987; Kaplan and Rossetti, 2011; Oddo et al., 2009). Unfortunately, most of the discussed papers limit their spectrum analysis to only one vigilance state, or in some cases to a single frequency band of interest. Where oscillatory activity is discussed, spectra were typically not decomposed into periodic and aperiodic components. Our results suggest, however, that oscillations are a main contributor to sepsis-related changes in power spectra in our model, while background shifts in E-I balance, as indicated by spectrum slope (Richard Gao et al., 2017), were less prominent. Further studies that consider both sleep and wake states will be required to further elucidate NREM-REM interactions and the role of oscillatory and aperiodic brain activity in sepsis.

One hypothesized cause of increased state switching is state instability, as evidenced by high epoch-to-epoch variability within the states, resulting in high state-space velocity values. In line with this, we found higher within-state velocities in the higher frequency spectral ratio (R2) after LPS injection. In septic rats, we found increased velocities in the higher frequency ratio, and not in the lower frequencies where we observed the largest overall shifts in cluster location. Changes in cortical higher frequency phenomena like spindles and HFOs signal NREM-REM transitions in normal sleep (Sánchez-López et al., 2018). Increased variability in higher EEG frequencies, even outside of the transition periods themselves, may help drive increased state switching in our animals as well. The relation between within-state velocity and sleep state stability is not entirely straightforward. Increased within-state velocities were previously observed in orexin knockout mice, but not in narcolepsy patientsdespite high degrees of state fragmentation in both (Diniz Behn et al., 2010; Schoch et al., 2017). In Parkinson’s disease, which is characterized by abnormally low velocities, lower within-state velocities were associated with decreased arousability rather than changes in gross sleep architecture (Imbach et al., 2016). Even so, higher within-state velocities seem to be generally related to instability, although this does not always rise to the level of increased vigilance state changes in these differing pathologies. Additionally, we found that in our sepsis model, the state transitions themselves remained normal. State-space parameters like cluster distance, velocity and transition quality can thus each be affected differentially according to underlying pathophysiology. Further research will be needed to elucidate mechanisms underlying these changes in acute inflammatory conditions like sepsis.

We performed recordings under urethane anesthesia, which is known to largely spare the ongoing brain oscillations and to mimic a number of physiological features of natural NREM and REM sleep (Clement et al., 2008; Pagliardini et al., 2013). This offers the possibility to analyse their fine kinetics in the absence of many peripheral inputs, pain, and awakening responses that complicate the study of sleep-wake-like rhythms in ICU settings and in non-anaesthetized animals. Our reduced complexity model captures many of the features of sepsis-associated changes in brain activity, but not all. Acute suppression of REM sleep is a common effect of inflammation under non-anaesthetized conditions (Opp, 2005), in addition to fragmentation of the remaining NREM sleep and wakefulness. Although we do not find lower amounts of the REM-like urethane state in the current study, we do find severe state fragmentation and spectral changes observed in other studies. Given the role of sleep in maintaining a functional immune system and its association to memory consolidation, it’s crucial to better understand sleep disturbances during sepsis. There is substantial proof of memory impairments after sepsis, but the role of disturbed sleep remains to be fully explored. Remedying sleep fragmentation could be a potential treatment option for eventual recovery of memory impairments and cognitive function in sepsis survivors. Here, we aimed at getting a step closer to understanding these sleep state dynamics.

## 5. Acknowledgements

The study was supported by Charles University projects START/MED/107, Cooperatio / NEUR and by project CZ.02.1.01/0.0/0.0/16_019/0000787 „Fighting INfectious Diseases”, awarded by the MEYS CR and financed from EFRR.

We thank to Lenka Sykorova and Stepan Kapl for technical support and to Karel Blahna for his valuable inputs.

## Abbreviations

EEG: electroencephalogram
LFP: local field potential
LPS: lipopolysaccharide
NREM: non-rapid eye movement sleep
REM: rapid eye movement sleep
SABD: sepsis-associated brain dysfunction
rmANOVA: repeated measures analysis of variance

