## supplemental figures for "Sepsis-induced brain state instability"

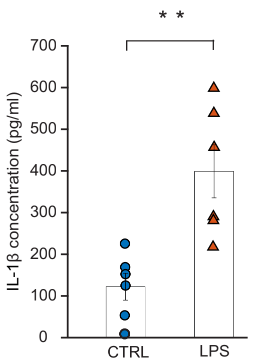


**Supplemental Figure S1.** LPS injection increased inflammatory marker IL -1β in LPS group post LPS injection.

Each point represents the blood concentration of IL -1β (pg/ml) in control or LPS rats 8-9 hours after saline or LPS injection. Bar graphs show mean ± s.e.m values of serum IL-1 β. **$p < 0.0$

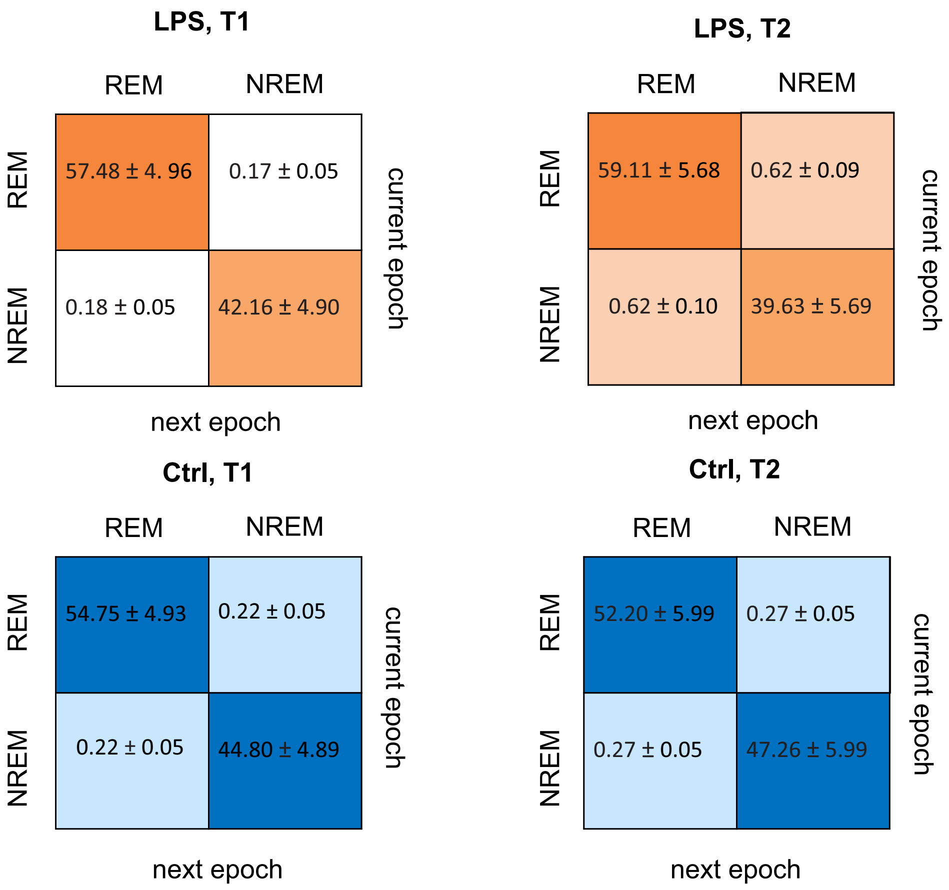


**Supplemental Figure S2.** LPS leads to increased transition probability from REM to NREM or NREM to REM.

Each cell shows average transition probability per epoch from REM to REM, REM to NREM, NREM to REM or NREM to NREM states for: Control, T1 (before injection), Control, T2 (after injection), LPS, T1 (before injection) and LPS, T2 (after injection).

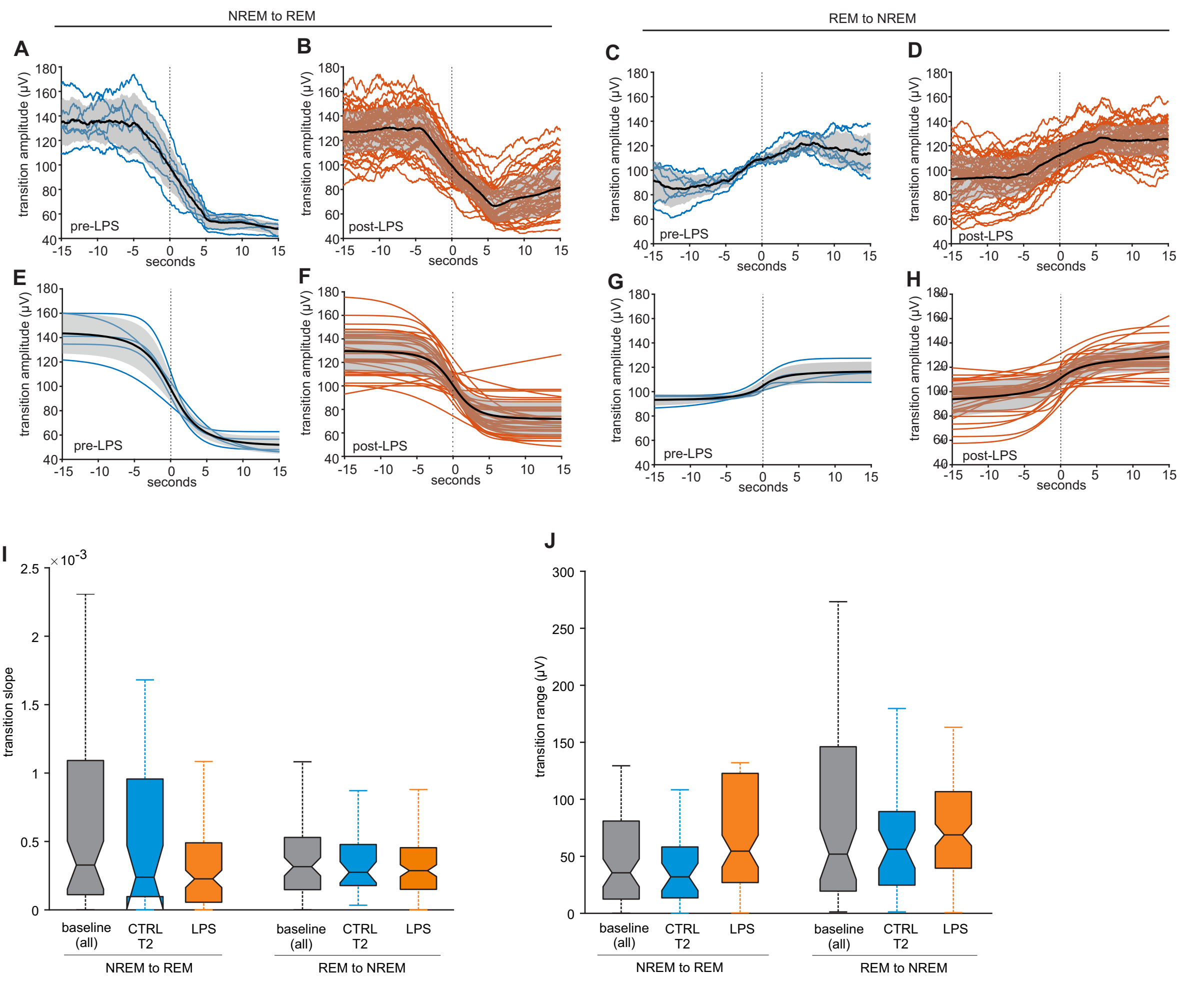


**Supplemental Figure S3.**
A-D: Smoothed EEG traces for state transitions for a representative animal at baseline (A,C) and after LPS injection (B,D). NREM-to-REM transitions are shown in A and B, REM-to-NREM transitions are shown in B and D. Mean traces are shown in black, with standard deviation in gray.

E-H: fitted curves for NREM-to-REM (E,F) and REM-to-NREM transitions (G,H) at baseline (E,G) and after LPS injection (F,H) for the traces shown in A-D. Mean fitted curves are shown in black, with standard deviation in gray.

I: Box plots showing medians and quartiles for maximum transition slope for all NREM-to-REM and REM-to-NREM transitions in all baseline recordings (N = 80 and N = 79, resp., 12 rats), T2 in controls (N = 37 and N = 34, resp., 6 rats), and after LPS injection (N = 123 and N = 116, resp., 6 rats).

J: Box plots showing medians and quartiles for transition range for all NREM-to-REM and REM-to-NREM transitions in all baseline recordings (N = 80 and N = 79, resp., 12 rats), T2 in controls (N = 37 and N = 34, resp., 6 rats), and after LPS injection (N = 123 and N = 116, resp., 6 rats).

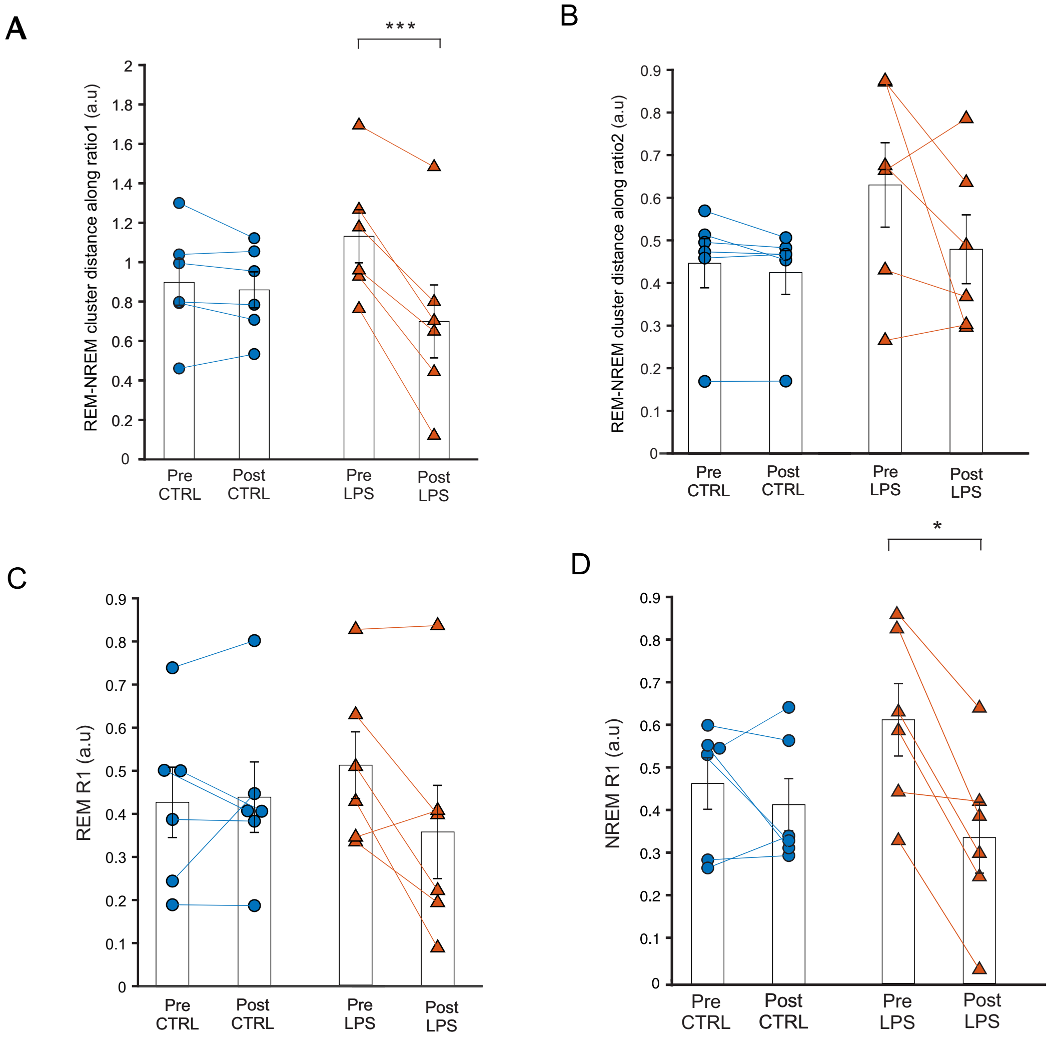


**Supplemental Figure S4.** LPS leads to spectral changes in the 1-9 Hz frequency range in both REM and NREM.

A, B Each data point represents the distance between REM and NREM cluster medians along ratio1 (A) or ratio 2 (B) in CTRL (circles) and LPS (triangles) animals pre or post injection. Bars represent the mean ± s.e.m. ***p<0.001

C, D Each data point represents the median of R1 of REM (C) or NREM (D) in CTRL (circles) and LPS (triangles) animals' pre or post injection. Bars represent the mean ± s.e.m.  **p<0.05; ***p<0.001.

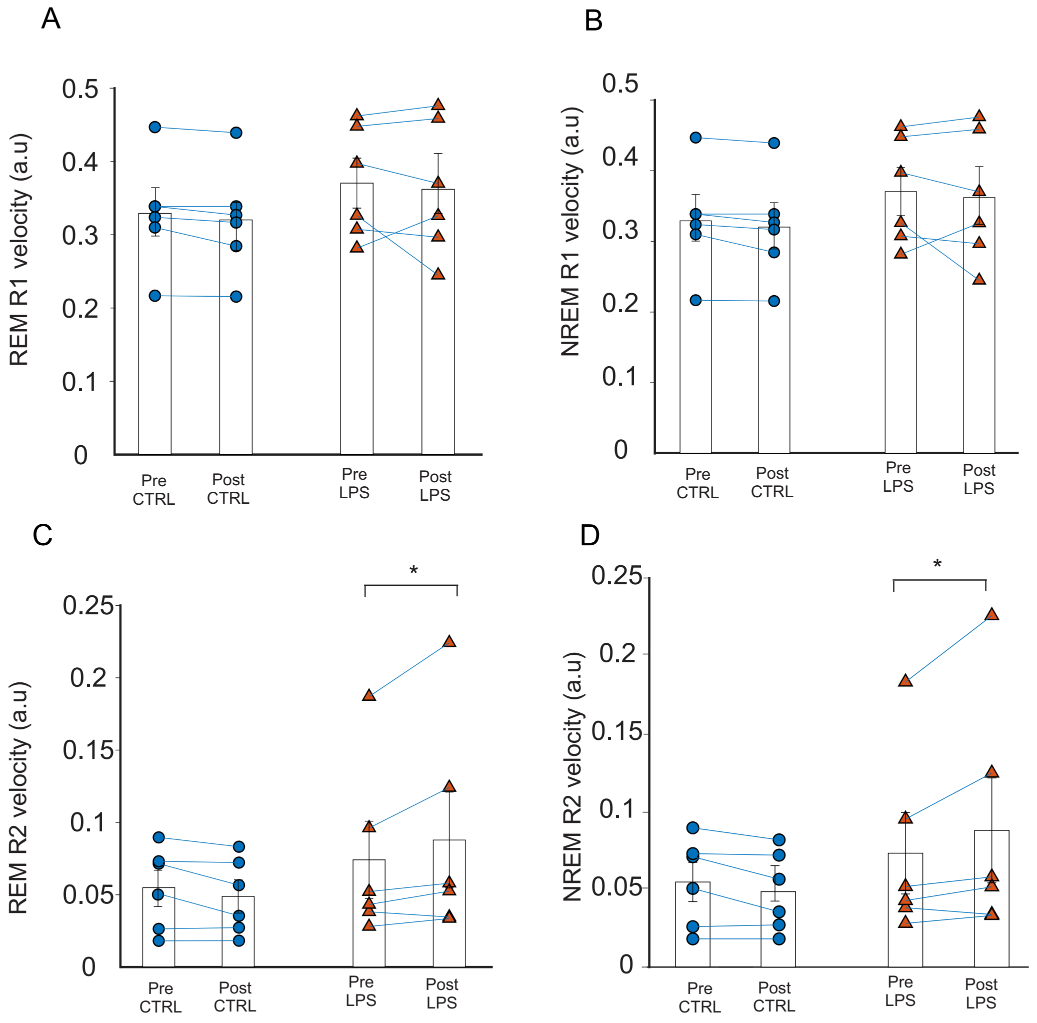


**Supplemental Figure S5.** LPS leads to sleep state instability in high frequency range, R2 (1-15 Hz/1-45 Hz).
A, B Median velocity of REM (A) and NREM (B) along R1 in CTRL (circles) and LPS (triangles) animals pre or post injection. Bars represent mean ± SEM.
C, D Median velocity of REM (A) or NREM (B) along R2 in CTRL (circles) and LPS (triangles) animals pre or post injection. Bars represent mean ± SEM. *p<0.05

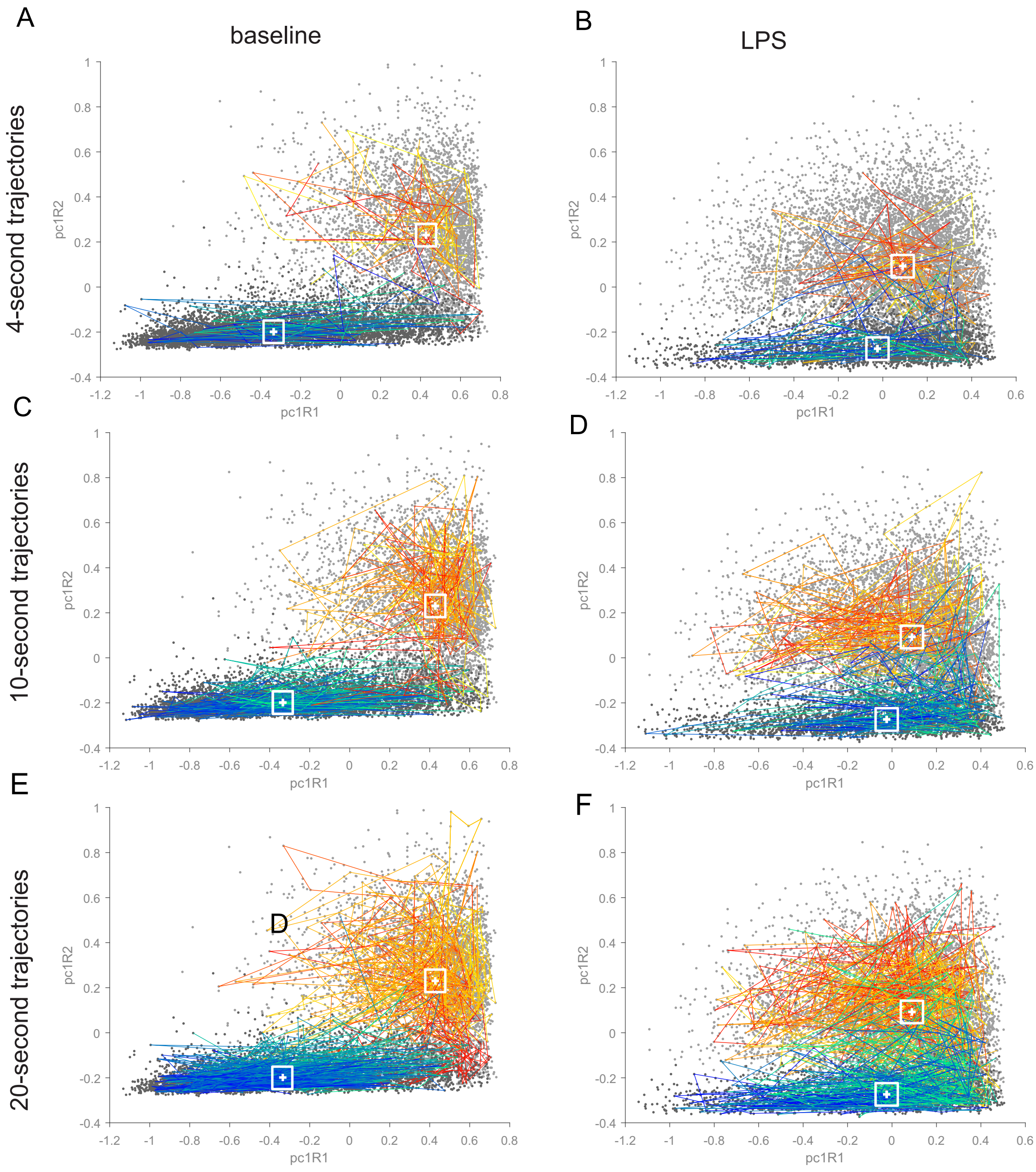


**Supplemental Figure S6.**Representative 4-, 10-, and 20- second long trajectories in state space. White boxes indicate the region around the state cluster median where the detected incoming and outgoing trajectories end resp. start.
15 incoming and 15 outgoing trajectories are shown for each state. Plotted trajectories were chosen at random.
Although second-to-second positions in state space vary widely (points that are neighboring in time are not necessarily closely spaced together in state space), the overall trajectories sampled around a cluster median remain mostly within the boundaries of that cluster. This is the case at baseline, but also after LPS injection. Similar patterns were observed for short trajectories (4s) and longer trajectories (20s).

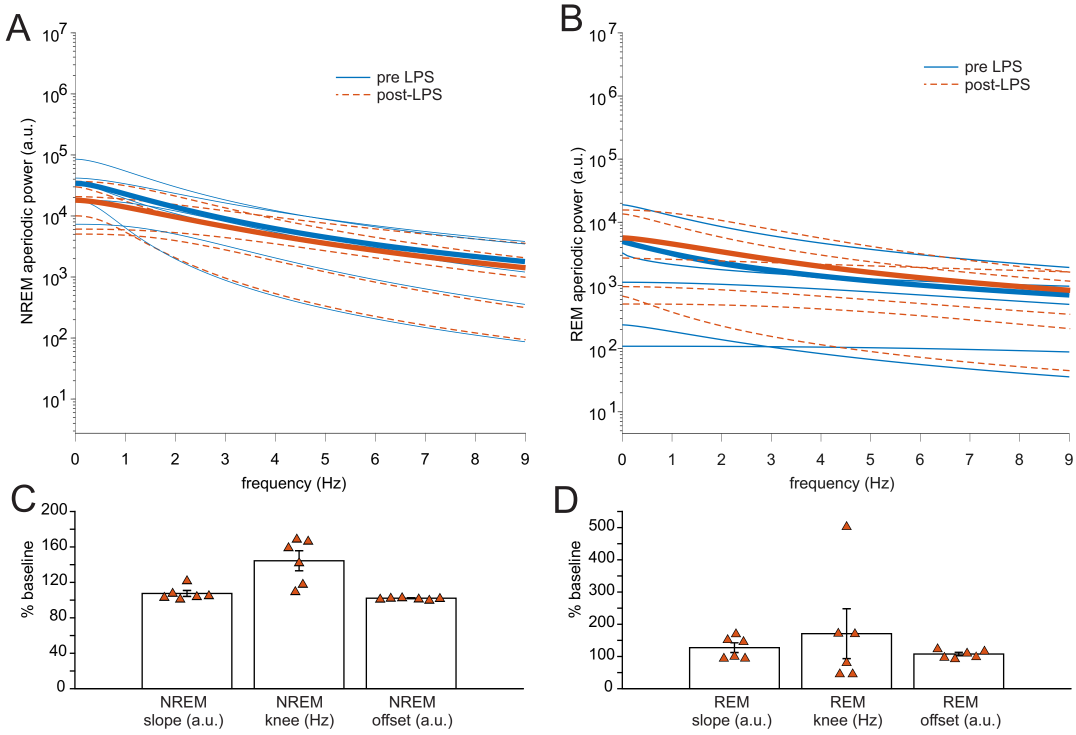


**Supplemental Figure S7.** Effects of LPS injection on aperiodic spectrum components

Aperiodic component of the FOOOF model in the NREM-like state (A) and the REM-like state (B) before (blue, solid lines) and after (orange, dashed lines) LPS injection. Thin lines show aperiodic components of individual spectra, thick lines show group average.

Post-LPS changes in individual variables determining the aperiodic spectral component: slope, knee frequency and offset in NREM (C) and REM (D).
Bars and error bars show mean + s.e.m, triangles show values for individual rats.
